# A second generation capture panel for cost-effective sequencing of genome regulatory regions in wheat and relatives

**DOI:** 10.1101/2022.09.16.508311

**Authors:** Junli Zhang, Juan M. Debernardi, German Burguener, Frédéric Choulet, Etienne Paux, Lauren O’Connor, Jacob Enk, Jorge Dubcovsky

**Affiliations:** University of California, Davis, CA 95616, USA; UCA, INRAE, GDEC, 63000, Clermont-Ferrand, France; VetAgro Sup, Lempdes, France; formerly INRAE; Daicel Arbor Biosciences, Ann Arbor, MI 48103, USA; Howard Hughes Medical Institute, Chevy Chase, MD 20815, USA

## Abstract

As genome resources for wheat expand at a rapid pace, it is important to update targeted sequencing tools to incorporate improved sequence assemblies and regions of previously unknown significance. Here, we developed an updated regulatory region enrichment capture for wheat (*Triticum* L.) and other *Triticeae* species. The core target space includes sequences from two Kbp upstream of each gene predicted in the Chinese Spring wheat genome (IWGSC RefSeq Annotation v1.1) and regions of open chromatin identified with ATAC-seq from wheat leaf and root samples. To improve specificity, we aggressively filtered candidate repetitive sequences using a combination of BLASTN searches to the Triticeae Repetitive Sequence Database (TREP), identification of regions with read over-coverage from previous target enrichment experiments, and k-mer frequency analyses. The final design comprises 216 Mbp of predicted hybridization space in hexaploid wheat and showed increased specificity and coverage of targeted sequences relative to previous protocols. Test captures on hexaploid and tetraploid wheat and other diploid cereals show that the assay has broad potential utility for cost-effective promoter and open chromatin resequencing and general-purpose genotyping of various *Triticeae* species.

## 1 INTRODUCTION

The coordinated action of promoters and enhancers is critical to regulate the precise spatial and temporal patterns of gene expression required for organismal development and appropriate responses to environmental changes (Wittkopp and Kalay, 2012). Sequence-level variation in regulatory elements is a major driver of phenotypic variation and adaptation (Rodgers-Melnick, Vera, et al., 2016) and, therefore, genomic tools are necessary to access this variability. This is particularly relevant in crop species, where variation in regulatory regions can be used to improve economically valuable traits. Variants in regulatory regions are especially useful for plant improvement projects because they usually cause less severe phenotypic changes than gene knockout mutations, since they are expected to affect the timing, spatial distribution or levels of gene expression rather than the function of all the transcripts (Rodriguez-Leal, Lemmon, et al., 2017, Wittkopp and Kalay, 2012).

In species with small genomes, such as *Arabidopsis* and rice (< 1 Gbp), natural variation in regulatory and coding regions can be cost-effectively accessed by whole genome resequencing of a large number of accessions. However, this strategy is not economically viable in polyploid species such as wheat, that have multiple copies of 4-5 Gb-sized subgenomes composed mainly of repetitive elements (International Wheat Genome Sequencing Consortium, 2018). Instead, capture platforms have been used to sequence the coding and regulatory regions of the wheat genome. A variety of exome-targeting probe sets for wheat have been designed using various commercial platforms resulting in useful studies (Chen, Hegarty, et al., 2021, Dang, Zhang, et al., 2022, Gabay, Zhang, et al., 2021, Glenn, Zhang, et al., 2022, Krasileva, Vasquez-Gross, et al., 2017, Serra, Svacina, et al., 2021). However, some of those capture assays have been discontinued and would be expensive to reproduce as new custom assays for small or moderate sample numbers. Currently, only the Wheat Exome version 1 design offered by Daicel Arbor Biosciences (hereafter “Arbor”) is available as an off-the-shelf wheat probe set.

A global wheat promoter capture assay was developed after the release of the Chinese Spring RefSeq v1.0 sequence (International Wheat Genome Sequencing Consortium, 2018) covering two Kbp of sequence upstream of the start codon of all high-confidence annotated genes (Gardiner, Brabbs, et al., 2019). However, this design is among the ones that were recently removed from public availability. In this study, we present a new improved and expanded regulatory capture design developed in collaboration with Arbor, which they offer as a catalog product for broad community use. Our new design leverages data obtained from capture experiments using the previous promoter probe design, eliminates previously undetected repetitive regions, and expands the scope of regulatory targets by including open chromatin regions identified with ATAC-seq (Assay for Transposase-Accessible Chromatin using sequencing) (Buenrostro, Giresi, et al., 2013) data from wheat leaf protoplasts (Lu, McKenzie, et al., 2020) and roots (Debernardi, Burguener, et al., 2022). We evaluate the behavior of the probe set for enrichment efficiency and overall genomic coverage depth and breadth in several *Triticum*, *Aegilops*, and *Secale* samples.

## 2 MATERIALS AND METHODS

### 2.1 Capture design: promoter sequences

Target design began with the same 2-Kbp regions upstream of all the high-confidence wheat genes in Chinese Spring (CS) annotation RefSeq v1.1 that were used in the previous promoter capture design (Gardiner, Brabbs, et al., 2019), henceforth ‘Gardiner probe set’. This initial sequence included 110,788 distinct regions covering a total of 221.6 Mbp. We then excluded all the repetitive elements annotated in the wheat genome annotation RefSeq v1.1 (International Wheat Genome Sequencing Consortium, 2018). The subtraction of the corresponding BED files yielded a reduced sequence space of 168.8 Mbp (23.6% reduction).

We complemented this set of putative, single-copy hexaploid promoter regions with sequences from the tetraploid wheat Kronos (PI 576168, *Triticum turgidum* L. subsp. *durum* (Desf.) Husn.). These sequences were derived from 40,975 contigs of Kronos (33.4 Mbp) assembled from Kronos reads that did not map to CS (Krasileva, Vasquez-Gross, et al., 2017). We then aligned these contigs with the CS promoter space using BLASTN (less stringent than read mapping) and selected 13.5 Mbp of putative promoter regions, bringing the working target space to 182.4 Mbp. To further reduce sequence redundancy, we clustered the collection of contiguous sequences using a threshold of 99% identity with the software CD-HIT-EST v4.7 (Fu, Niu, et al., 2012), which resulted in 4.3% reduction of the sequence space to 174.6 Mbp. We then eliminated contiguous sequences smaller than 100 bp, bringing the final starting space for probe design to 174.0 Mbp represented by 167,685 contiguous sequences.

To eliminate repetitive sequences missed in the RefSeq v1.1 annotation, we performed additional filtering steps. First, we used BLASTN to query our 167.7 K target sequences against the database of Triticeae Repetitive elements (TREP) (Wicker, Matthews, et al., 2002) and removed significant similar sequences (E < 1e^−10^). Then, we performed a k-mer analysis using Tallymer v1.6.1 (Kurtz, Narechania, et al., 2008) and an index previously created based on Chinese Spring RefSeq v1.0 (k-mer length = 17) (International Wheat Genome Sequencing Consortium, 2018). We evaluated 4 different thresholds of k-mer occurrence: 5, 10, 50 and 100, which masked 55%, 36%, 13%, and 8% of the regulatory regions, respectively. To avoid masking conserved regulatory regions, we decided to mask 17-mers repeated 100 times or more in the genome. Finally, we analyzed twenty captures performed in tetraploid Kronos with the Gardiner probe set, and masked regions with a coverage higher than 100×, which is >5-fold higher than the average coverage of these captures. The additional masking generated new fragmentation, so we removed one last time sequences of contiguous length lower than 100 nt. The final targeted putative promoter space was 162.5 Mbp.

### 2.2 Capture design: open chromatin regions from ATAC-seq

To include additional regulatory sequences, we added open chromatin regions from publicly available wheat ATAC-seq data from wheat leaf protoplasts (Lu, McKenzie, et al., 2020) and from seminal roots recently generated in our lab (Debernardi, Burguener, et al., 2022). Root ATAC-seq data was generated from the tetraploid cultivar Kronos, while leaf ATAC-seq data was generated from leaf protoplast isolated from the hexaploid cultivar Paragon. Raw data from both studies was analyzed using MACS2 with the same parameters (Debernardi, Burguener, et al., 2022).

The initial ATAC-seq peaks covered 28.36 Mbp (57,981 peaks) in the leaf protoplast data and 2.75 Mb in the frozen root data (7,269 peaks). Among these peaks, we identified 4,432 (61%) that overlapped in at least 10% of their length, suggesting a substantial proportion of shared open chromatin regions. Excluding the 1.53 Mbp of overlapping sequences, we identified 29.58 Mbp of non-redundant ATAC-seq data between the two datasets. From this we subtracted the 6.1 Mbp already present in the promoter sequences, and added 23.5 Mbp of new sequences to the regulatory target design.

### 2.3 Probe design

Each selected target sequence from the promoters and ATAC-seq was padded by 100 nt on either side and re-merged, comprising 241.59 Mbp of potential hybridization target. We used an “island” approach for tiling probes, wherein we selected 80-nt probes with the best predicted hybridization dynamics in every 100-nt window across the padded space. These probe candidates were then aggressively filtered for specificity against both RefSeq v1.0 and the draft Kronos assembly in order to once again minimize the likelihood of targeting multi-copy sequence motifs. To describe the final hybridization target space on RefSeq 1.0, we mapped the probes to the reference genome using *bwa mem* version 0.7.10-r789, and padded these map sites by 100 nt to reflect a likely retrievable space of roughly 216.47 Mbp (DAB_WheatRegulatoryV1.IWGSCv1_hybspace.bed), which we refer to hereafter as “hybridization space”. Roughly 168.0 Mbp of this space intersects with the original non-padded target space on hexaploid RefSeq v1.0, which we refer to as the final “target space” (DAB_WheatRegulatoryV1.IWGSCv1.bed). To determine the corresponding regions on the rye assembly, we used BLAST blastn (version 2.6.0+) to map the target and capture space wheat sequences to the rye genome, after which the top bit score intervals per query sequence were retained and merged (DAB_WheatRegulatoryV1.Weiningv1.bed and DAB_WheatRegulatoryV1.Weiningv1_hybspace.bed, respectively).

The probe set was synthesized in subgenome-specific modules, so that a user can exclude probes for a subgenome not present in the sample being enriched, or otherwise customize the composition of enrichment reactions. Supplemental Table S1 summarizes the total target and hybridization spaces for each of the reference genomes and sub-genomes (for the polyploid species), as well as the names of the BED files deposited in GitHub (https://github.com/DubcovskyLab/DAB_WheatRegulatoryV1).

### 2.4 Capture performance evaluation

As test material, we first generated Illumina TruSeq-style sequencing libraries from genomic DNAs extracted from eight hexaploid accessions (*T. aestivum*, genomes ABD), 24 tetraploid Kronos EMS mutants (*T. turgidum* subsp. *durum*, genomes AB) (Krasileva, Vasquez-Gross, et al., 2017), and 16 diploid accessions. The diploid accessions included eight from *T. monococcum* (A^m^ genome), two from *T. urartu* (A genome), three from *Aegilops speltoides* (S genome), one from *Aegilops markgrafii* (C genome) and two from *Secale cereale* (R genome) (Supplemental Table S2).

Genomic DNA was sonicated with a Q800R instrument (Qsonica, Newtown, CT, USA) to mean lengths of 400 bp and purified with dual-sided SPRI treatment. Then either 200 ng (hexaploid and tetraploids) or 100 ng (diploids) sonicated and size-selected gDNA was taken to end repair, A-tailing, and adapter ligation using the KAPA HyperPrep DNA kit (Roche). Each ligation product was index-amplified using unique dual 8-bp indexing primers for eight cycles with KAPA HiFi polymerase (Roche).

For hexaploid *T. aestivum*, Kronos-specific probes were excluded from the captures. For tetraploid *T. turgidum* ssp. *durum* captures, the D-specific probes were excluded. For diploid wheat and rye, we captured several taxa in the same pools, so we included all probe modules. For the captures, we pooled 8 libraries in hexaploid wheat (1 μg each), 12 in tetraploid wheat (750 ng each) and 16 in the diploid species (500 ng each). Captures were conducted following the protocol described in myBaits Expert Wheat Exome kit (https://arborbiosci.com/wp-content/uploads/2021/08/myBaits_Expert_WheatExome_v1.51_Manual.pdf).

We sequenced the resulting capture pools on an Illumina NovaSeq 6000 S4 flow-cell using a PE150 protocol, with sample demultiplexing requiring 100% match to both the i5 and i7 indexes. To analyze target enrichment specificity, we down-sampled the reads to 1 M read-pairs (2 M reads) prior to reference mapping. For analysis of coverage depth and breadth and for variant calling, we down-sampled to 60 M pairs (18 Gbp) for the hexaploid accessions, 40 M pairs (12 Gbp) for the tetraploid accessions, and 20 M pairs (6 Gbp) for the diploid species, which follows recommended sequencing depths for analysis of data generated with Arbor’s Wheat Exome V1 kit. Some samples did not yield this minimum number of reads and so were excluded from coverage analysis (Supplemental Table S2). After down-sampling, reads were taken directly to reference alignment with *bwa mem* (version 0.7.10-r789) to either taxon-appropriate subgenome sets of Chinese Spring RefSeq v1.0 (International Wheat Genome Sequencing Consortium, 2018), or to the genome assembly of Weining rye (Li, Wang, et al., 2021). In both cases, mapping was performed to “parts” versions of each genome assembly, wherein each chromosome was divided into two (wheat) or three (rye) parts to make them compatible with *bwa* reference indexing. Following PCR duplicate removal with *picard MarkDuplicates* (version 2.18.15-SNAPSHOT), coverage depth and breadth were assessed with *bedtools* (version 2.17.0) and variants were called using *bcftools* (version 1.10.2-105-g7cd83b7, default parameters) with a minimum quality score of 20 and read depth of 10.

### 2.4 Comparison of this and previous promoter capture assays in tetraploid wheat

To compare the behavior of this new capture probe set and protocol to those described by Gardiner et al. (2019), we enriched libraries built from twenty-four Kronos EMS-mutagenized lines included in a previous exome capture study (Krasileva, Vasquez-Gross, et al., 2017). DNA extraction, construction of the sequencing libraries and capture followed procedures described previously (Krasileva, Vasquez-Gross, et al., 2017). Target enrichment was performed in the same set of 24 tetraploid samples using both the promoters-only probes in the Gardiner probe set, ordered as a custom kit from Roche (SeqCap EZ Prime Developer Probes, cat# 8247633001), and the Arbor assay described in this study.

Each enrichment reaction was performed on a pool of 12 libraries (125 ng per library to be consistent with previous protocols). Following enrichment, the captured DNA was amplified for ten cycles using KAPA HiFi HotStart ReadyMix (6.25 ml, Roche, catalog number 7958935001) and purified in 1.8 x volume of Agencourt AMPure beads (Beckman Coulter, catalog number A63881). Captured DNA was eluted in 30 μL of ultrapure water and quantified using QUBIT 2.0. Following enrichment, the pools were sequenced on one lane of Illumina NovaSeq S4 (PE150) at the Genome Center of UC Davis, and informatically processed in identical fashion as the main capture tests.

After sequencing, we used a similar analysis procedure as was used for general capture metrics in our test set. First, read data was down-sampled to the same level as before (40 M pairs). Then we estimated the target region coverage depth (as defined by the subgenome A, B, and Un subgenome entries of “Prom-capture-HC+5UTR-targets.bed” from Gardiner et al. 2019), the percent of reads on-target and the percent of duplicated reads to gauge overall library complexity at this raw read depth. We also compared the number of EMS mutations detected with the two capture protocols using the MAPS pipeline described previously (Henry, Nagalakshmi, et al., 2014). This pipeline eliminates polymorphisms between Kronos and Chinese Spring and potential homeologous polymorphism by excluding duplicated SNPs in sets of 24 libraries analyzed simultaneously (Henry, Nagalakshmi, et al., 2014, Krasileva, Vasquez-Gross, et al., 2017). To avoid sequencing errors, we only called mutations that were detected at least four times in heterozygous mutations and at least 3 times in homozygous mutations. In the previous exome capture, this threshold resulted in an error rate lower than 0.3% (Krasileva, Vasquez-Gross, et al., 2017).

## 3 RESULTS

### 3.1 Promoter Capture and ATAC-seq

The combined 2-kb promoter regions in front of all the high confidence annotated genes resulted in an initial 221.6 Mbp target space. However, after the multiple filtering steps for repetitive sequences described in the Materials and Methods section, this space was reduced to 162.5 Mbp (26.7% reduction). These promoter sequences were then complemented with open chromatin sequences obtained from roots and leaf protoplasts ATAC-seq data (Debernardi, Burguener, et al., 2022, Lu, McKenzie, et al., 2020).

Figure 1 shows an example of a good overlap between peaks from the leaf protoplast and the seminal root tips. It also shows the presence of open chromatin regions located outside the 2 Kbp regions upstream of the start codons selected for the first promoter capture. The combined analysis of the open chromatin regions revealed that approximately 75% were outside the promoter regions (Debernardi, Burguener, et al., 2022), indicating that a large proportion of potential regulatory elements can be missed in captures based on promoter sequences only. This emphasizes the importance of complementing promoter regions with open chromatin data to maximize the detection of variation in regulatory regions.

**Figure 1.**
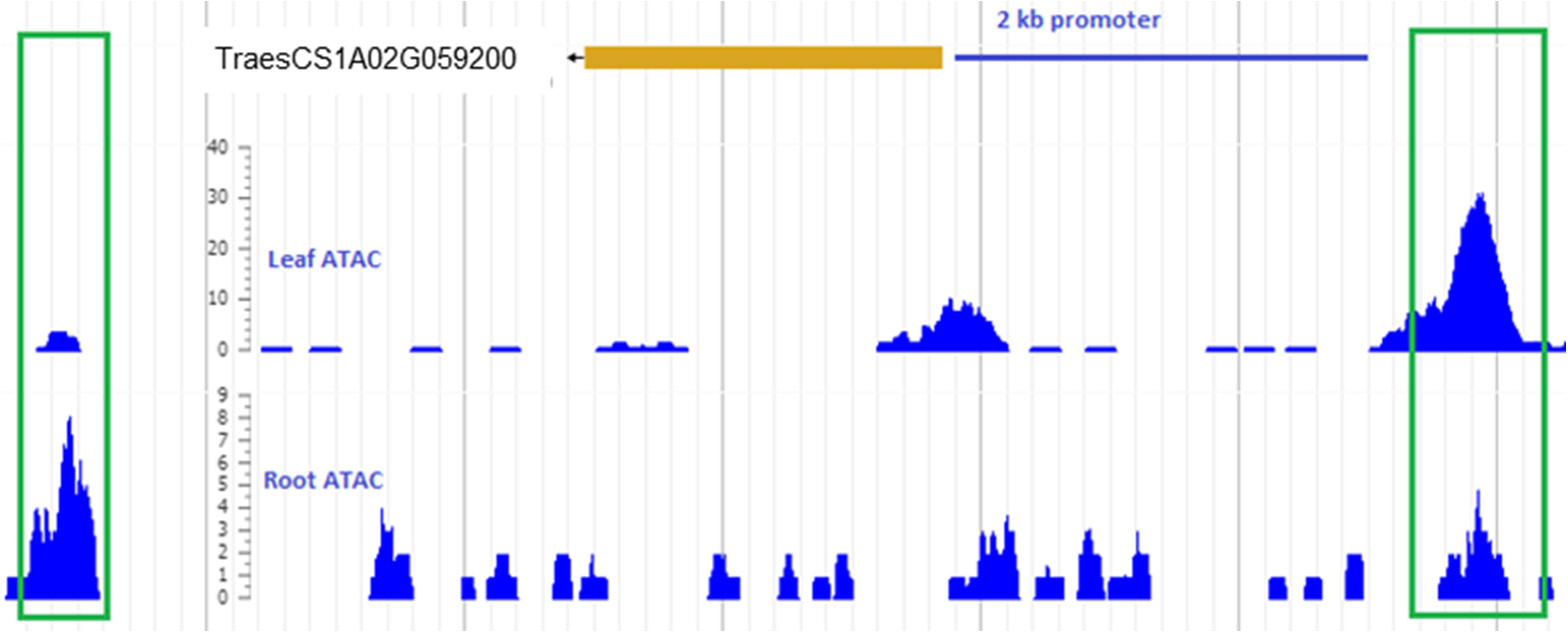
Example of open chromatin regions detected in the leaf protoplast and seminal root tips by ATAC-seq. The green rectangles indicate open chromatin regions detected outside the 2 kb promoter region.

The combined promoter and ATAC-seq data resulted in a final target space of 186.0 Mbp. After padding each selected target with 100 nt on either side the final hybridization target space was 241.59 Mbp. The target and hybridization spaces by genome are described in Supplemental Table S1.

### 3.2 Capture performance

Capture performance was evaluated for specificity (i.e., non-target exclusion or “reads on-target”) as well as for coverage breadth and depth. To ensure a fair coverage evaluation between libraries, we sampled identical numbers of read pairs based on sample taxon ploidy, and then aligned the reads to their most closely-related RefSeq or Weining Rye (sub)genomes (Figure 2a). Our sampling requirements did eliminate seven of the 48 sequenced libraries from coverage analysis but all taxa were represented by at least one library with sufficient data, and in most cases several (Supplemental Table S2).

**Figure 2.**
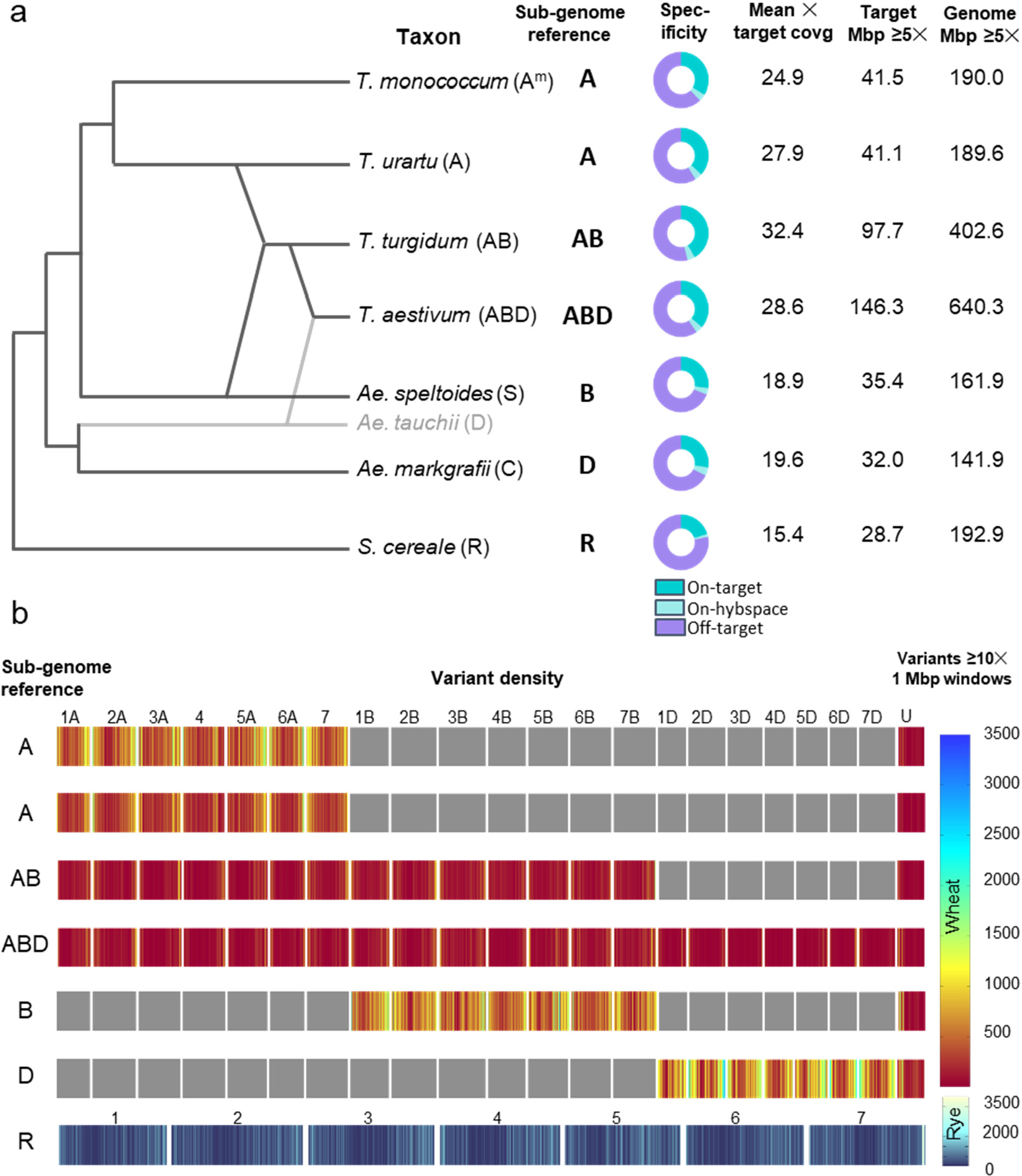
Performance of the new regulatory region capture assay. (a) target coverage depth and genome coverage breadth observed in diploid, tetraploid and hexaploid wheat accessions and rye. Genome designations are indicated in parenthesis after the species name followed by the sub-genomes used as reference for mapping. For diploid libraries, raw read data was down-sampled to 20 M read-pairs, tetraploid to 40 M, and hexaploid to 60 M. Alignment was performed on either the entire Chinese Spring RefSeq v1.0 genome reference, or on the subgenome(s) appropriate for the taxon. Specificity is measured as the proportion of raw reads overlapping the target (blue) or predicted hybridization space (light blue) in the circular graphs. Raw data is presented in Supplementary Tables S2 and S3. Coverage depth and breadth were measured on the target space (~168 Mbp in hexaploid wheat) and on the entire genome after PCR deduplication and merging of overlapping R1-R2 pairs. (b) SNP distribution on different chromosomes. SNPs were called for sites with a minimum quality score of 20 and minimum depth of 10 unique reads.

Results are summarized in Supplemental Table S2 and averages per species presented in Supplemental Table S3 and Figure 2a. Specificity of the assay was measured by counting the proportion of non-deduplicated reads or read-pairs that overlapped either the target or hybridization space. Coverage depth and breadth of the target and genome overall was measured following PCR duplicate collapse and consolidation of overlapping R1 and R2 read-pair coordinates to single intervals. Finally, to compare repeat content retrieved with this and the Gardiner probe set captures, we measured the number of raw reads that mapped to unique locations in the genome.

The average specificity of the eight hexaploid libraries (one 8-plex capture) varied depending on the method used between 36.3% on-target to 40.5% on-hybridization space for single reads, and from 39.9% on target to 43.0% on-hybridization-space for read-pairs (Supplemental Table S3). Tetraploid captures (two 12-plex captures) were on average more specific (41.7% reads on-target to 49.8% read-pairs on-hybridization-space). In the diploid species, specificity decreased with increased divergence from the genome used as reference in the assay. *T. urartu*, which is the donor of the A genome in polyploids wheat, showed the highest specificity (36.8% reads on-target to 46.6% read-pairs on-hybridization-space, Supplemental Table S3), followed by *T. monococcum* (33.7% reads on-target to 43.2% read-pairs on-hybridization space) which diverged from *T. urartu* one million years ago (Dubcovsky and Dvorak, 2007). Specificity was further reduced in the more distantly related *Ae. speltoides* (27.1% reads on-target to 36.3% read-pairs on-hybridization-space) and *Ae. markgrafii* (27.6% reads on-target to 37.5% read-pairs on-hybridization-space), and the lowest specificity was observed for *S. cereale* genome (20.1% reads on-target to 27.4% read-pairs on-hybridization-space). In summary, specificity showed a good correlation with the evolutionary history of these species (Figure 2a, Supplemental Table S3).

Per-base mean coverage of the hybridization-space (Supplemental Table S3) was highly correlated with assay specificity (*R* = 0.9871, *P* < 0.0001). Tetraploid and hexaploid accessions exhibited on average 29.4 × and 26.1 × hybridization-space coverage. These values then decreased with phylogenetic distance to 25.4 × in *T. urartu*, 22.6 × in *T. monococcum*, 18.1 × in *Ae. markgrafii*, 17.2 × in *Ae. speltoides*, and 14.4 × in rye (Supplemental Table S3). The “Target Region with at Least 5× Coverage” corresponds closely with ploidy level, but within ploidy level it was also affected by specificity and coverage (Supplemental Table S3). This region was 41.1 and 41.5 Mbp in the related diploid species *T. urartu* and *T. monococcum*, respectively, which is approximately half of the space in tetraploid wheat (97.7 Mbp) and one third of the space in hexaploid wheat (146.3 Mbp). Within the more distantly related diploid species this space decreased to 35.4 Mbp in *Ae. speltoides*, 32.0 Mbp in *Ae. markgrafii*, and 28.7 Mbp in rye (Supplemental Table S3).

Remaining non-target reads yielded broad general coverage of the genome. With this assay, we obtained an average genome space covered at 5× or higher of ~640.3 Mbp in hexaploid wheat, 402.6 Mbp in tetraploid wheat, and 141.9 Mbp to 192.9 Mbp in the other species. This shows that while the high-coverage regions are dominated by the intended promoter and open chromatin targeted regions, the assay does yield substantial genome-wide information that could be useful for a range of study applications, particularly those aimed at characterizing natural variation and SNP discovery.

Called variant densities are depicted in Figure 2b for one representative sample for each taxon and for sites with a minimum of 10 × unique read coverage. SNP densities and distribution are broadly consistent with the genome biology and known level of sequence divergence between the different tested taxa and the hexaploid reference genome. For instance, the A^m^ genome from *T. monococcum* and the B-like genome of *T. speltoides* show the highest density of variants and the D genome of hexaploid wheat shows the lowest. In the wheat genome, genes and their regulatory elements are concentrated in the distal regions, and this is recapitulated in the variant density plots. The lower frequency of variants close to the centromeric regions is evident in several of the chromosomes. Rye is an outcrossing species with high levels of polymorphism, which is also reflected in the distribution of variants along its seven chromosomes. This high level of polymorphism may help explain the highly divergent performance metrics of the two rye specimens analyzed here, both in terms of read mappability to the genome, as well as specificity. In contrast, for the wheat species, overall performance was highly consistent both among different representatives of the same taxon, and among replicate libraries from the same genomic DNA source.

In summary, this capture assay was very efficient to capture variation in the regulatory regions of wheat and its closely related species, but has a lower specificity and covered a smaller target space when used in more distantly related species such as rye.

### 3.3 Comparisons of regulatory capture assays

The same twenty-four tetraploid wheat genomic DNA samples that were library-prepped and captured with this new regulatory design were also separately library-prepped and captured using a previous promoters-specific probe design (Gardiner et al. 2019) and the SeqCap EZ platform (Roche). To make an even comparison, we down-sampled to 40 M read-pairs (80 M reads) per library from both experiments. We mapped the reads to the Chinese Spring RefSeq v1.0 and identified EMS mutations using the MAPS pipeline (see Materials and Methods). We discovered an average of 3,338.6 and 3,131.5 EMS mutations per line with the Arbor and Gardiner probe sets and protocols, respectively (6.6% increase, *P* < 0.001, Table 1 and Supplemental Table S4). Both captures showed good proportion of mapped reads and similar proportion of G to A or C to T mutations, which are typically generated by EMS.

**Table 1.**
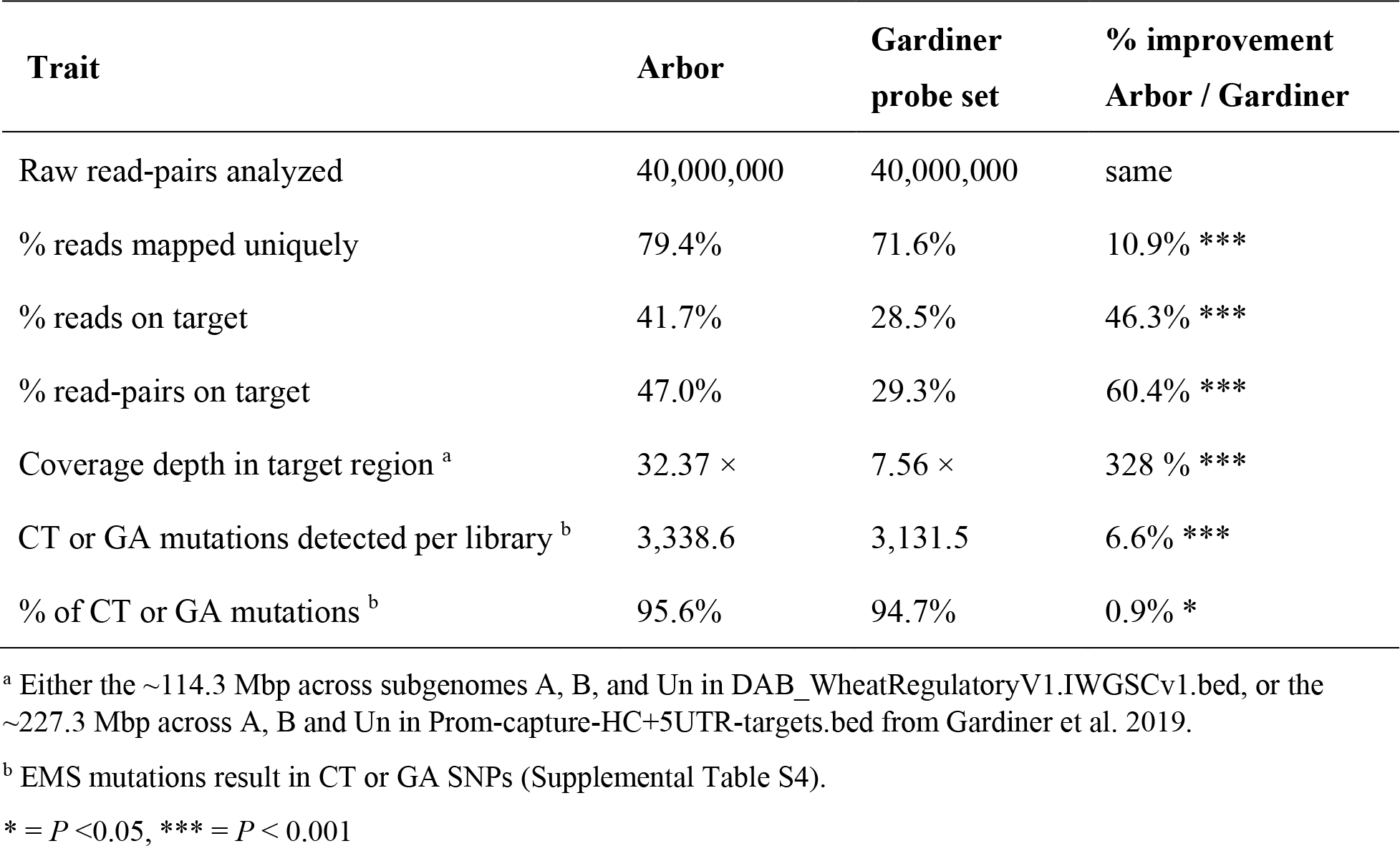
Comparison between the first and new wheat regulatory captures. Data are presented as the mean statistics of all 24 analyzed libraries.

Compared to the Gardiner probe set and protocol, the Arbor assay and protocol showed a significant (*P* < 0.001) increase in specificity (46.3% reads and 60.5% read-pairs on target) and coverage in the target region (3.3-fold increase, Table 1). The total number of reads mapping to the genome uniquely was 10.9% higher for the Arbor system, which indicates a higher rate of reads derived from putative single-copy loci. This particular metric should be robust to variation in library preparation between protocols, though differences in pre- and post-capture PCR amplification could contribute to the differences.

Taken together, these results suggest that the more stringent filtering of repetitive regions may have contributed to a significant reduction in duplicated reads and an increase of reads on target in the new protocol relative to the original one, although it is also possible that the different protocols contributed to these differences. Regardless of the cause of the differences, the new design and capture system described here represents an improvement in overall performance compared to our trials of the previous capture product.

## 4 DISCUSSION

The same myBaits technology used for this new regulatory capture design was used successfully previously in a more limited analysis of a subset of wheat gene promoters (Hammond-Kosack, King, et al., 2021). That study explored the 1.7 Kbp upstream of the coding regions of 459 wheat genes associated with agriculturally important traits in 95 ancestral and commercial wheat accessions. The study revealed a high level of conservation in the wheat promoter regions but also discovered many SNPs and indels located within predicted plant transcription factor binding sites. The new myBaits assay developed here expands the analyzed promoter regions by 200-fold to the promoters of all high confidence genes in the CS RefSeq v1.1 annotation.

A key special feature of this new capture assay is that it targets 23.5 Mbp of open chromatin detected with ATAC-seq data generated from leaf protoplasts (Lu, McKenzie, et al., 2020) and seminal root tips (Debernardi, Burguener, et al., 2022). Since transcription factors require open chromatin to exert their regulatory functions, the identified ATAC-seq regions present an excellent tool for identifying potential regulatory regions in the genome. Our analysis of the root ATAC-seq data showed a distribution of peaks among genic, promoter and intergenic regions that was similar to ATAC-seq data reported from other plant species (Maher, Bajic, et al., 2018). This open chromatin distribution indicates that a large proportion of putative regulatory regions can be missed by focusing only in the 2 Kbp regions upstream the start codons.

We observed a significant overlap (60%) between the ATAC-seq peaks detected in the leaf protoplast and the seminal root tips, in spite of the divergent nature of the tissues and conditions investigated. This level of overlap is comparable with the 71% overlap detected between ATAC-seq data from the more related root hairs and non-hair root cells reported in Arabidopsis (Maher, Bajic, et al., 2018). In addition, more than 99% of the peaks detected in the root frozen tissues were validated in an independent ATAC-seq study using fresh root tissues that yielded 7-fold more peaks (Debernardi, Burguener, et al., 2022).

The presence of a substantial number of tissue specific peaks (~40%) suggests that a more complete inventory of open chromatin regions will require additional ATAC-seq data from different wheat tissues, developmental stages and stress conditions. Therefore, additional capture assays will be necessary in the future to include these additional regulatory sequences.

## Abbreviations

ATAC-seq: Assay for Transposase-Accessible Chromatin using sequencing
BED: Browser Extensible Data
IWGSC: International Wheat Genome Sequencing Consortium
MAPS: mutations and polymorphisms surveyor
TREP: Triticeae Repetitive Sequence Database

## ACKNOWLEDGEMENTS

We thank Dr. Antony Hall (Earlham Institute, UK) for providing the sequences of the 2 Kbp regions in front of all the high confidence genes in the CS RefSeq v1.1 annotation, and Hans Vasquez-Gross for his help with the capture design. We also thank Jonathan Jones and Sebastian Fairhead of the Sainsbury Institute, UK, for contributing test genomic DNA. Finally, we thank the International Wheat Genome Sequencing Consortium (IWGSC) for their help to coordinate and disseminate this project. This project was supported by Agriculture and Food Research Initiative Competitive Grant 2022-68013-36439 (WheatCAP) from the USDA National Institute of Food and Agriculture.

## DATA AVAILABILITY STATEMENT

Data is publicly available. The names of the BED files for Target space wheat (DAB_WheatRegulatoryV1.IWGSCv1.bed.gz), hybridization space wheat (DAB_WheatRegulatoryV1.IWGSCv1_hybspace.bed.gz), Target space rye (DAB_WheatRegulatoryV1.Weiningv1.bed.gz), and hybridization space rye (DAB_WheatRegulatoryV1.Weiningv1_hybspace.bed.gz) are deposited in GitHub (https://github.com/DubcovskyLab/DAB_WheatRegulatoryV1). All Kronos mutant lines are available upon request from UC Davis and from the John Innes Center in the UK. The capture Assay is commercially available form Arbor Biosciences.

## AUTHOR CONTRIBUTIONS STATEMENT

JZ and LO performed all laboratory work. JMD performed the ATAC-seq of roots, JD and GB selected target regions. FC and EP contributed to target space filtration. JE designed the probe set. GB, JZ and JE performed bioinformatics analyses of experimental data. JZ and JD drafted the first version and all authors revised the manuscript. JD and JE designed, supervised, and secured funding for the project.

